# Microbial biosensor for sensing and treatment of intestinal inflammation

**DOI:** 10.1101/2023.07.21.550106

**Authors:** Duolong Zhu, Jeffrey Galley, Jason Pizzini, Elena Musteata, Jeffrey J. Tabor, Robert A. Britton

## Abstract

Substantial synthetic biology efforts have been made to engineer biosensors to detect intestinal inflammation, however none target the most clinically accepted biomarker, calprotectin. To develop an *in situ* biosensor for calprotectin, we optimized a zinc uptake regulator (Zur) regulated promoter coupled with a memory circuit that can detect and record intestinal inflammation *in vivo*. The level of activation strongly correlates with calprotectin levels in the colon of two independent mouse models of colitis. Coupling of the biosensor with the production of the anti-inflammatory cytokine IL-10 allowed for the resolution of chemically induced colitis, demonstrating the ability of the biosensor to sense and respond to disease. This work highlights the utility of developing synthetic organisms for the diagnosis and treatment of intestinal disease using clinically validated biomarkers.

**One sentence summary:** We have optimized a microbial biosensor to detect and respond to the clinically relevant intestinal inflammation biomarker calprotectin.

## Introduction

The use of engineered microbes to sense and respond to disease states is a central goal of synthetic biology (*1-3*). The intestine is a logical choice for employing engineered microbes as biosensors and for drug delivery as microbes have had a long evolutionary interaction with the human gut. Bacteria that can function as inflammation biosensors in animal models have been described (*1, 4, 5*). These biosensors target thiosulfate, tetrathionate, pH and reactive oxygen species and are based on observations made in mouse models of intestinal damage (*6-8*). However, none of these compounds are biomarkers for human disease and their utility to function in humans remains to be tested.

Monitoring of intestinal inflammation in patients in remission for the inflammatory bowel diseases Ulcerative Colitis and Crohn’s disease is challenging since the most widely accepted non-invasive test for measuring inflammation is based on fecal concentrations of the neutrophil protein calprotectin (*9, 10*). Calprotectin, which composes ∼50% of the effector molecules released by neutrophils upon degranulation during the antigenic response, is a heterodimer composed of the S100A8 and S100A9 Ca^2+^ binding peptides (*11, 12*). This protein sequesters essential metals (Zn^2+^ > Fe^2+^ > Mn^2+^) in the inflamed environment to restrict bacterial growth in a mechanism referred to as nutritional immunity (*13-15*).

We postulated that the human probiotic bacterium *Escherichia coli* Nissle (hereafter referred to as EcN) would be an ideal strain to develop an inflammation biosensor since it thrives in the inflamed gut, has had to evolve mechanisms for acquiring metals for growth in the presence of neutrophil infiltration, and is amenable to precision genome engineering (*16, 17*). Thus, we engineered calprotectin-responsive microbial biosensors that are capable of detecting and recording gut inflammation and *in situ* secreting therapeutic recombinant human IL-10 carried by YebF (secIL10) (*18*) to ameliorate intestinal inflammation (Fig. 1A). This work highlights the utility of developing synthetic organisms for the diagnosis and treatment of intestinal disease using clinically validated biomarkers.

**Fig. 1.**
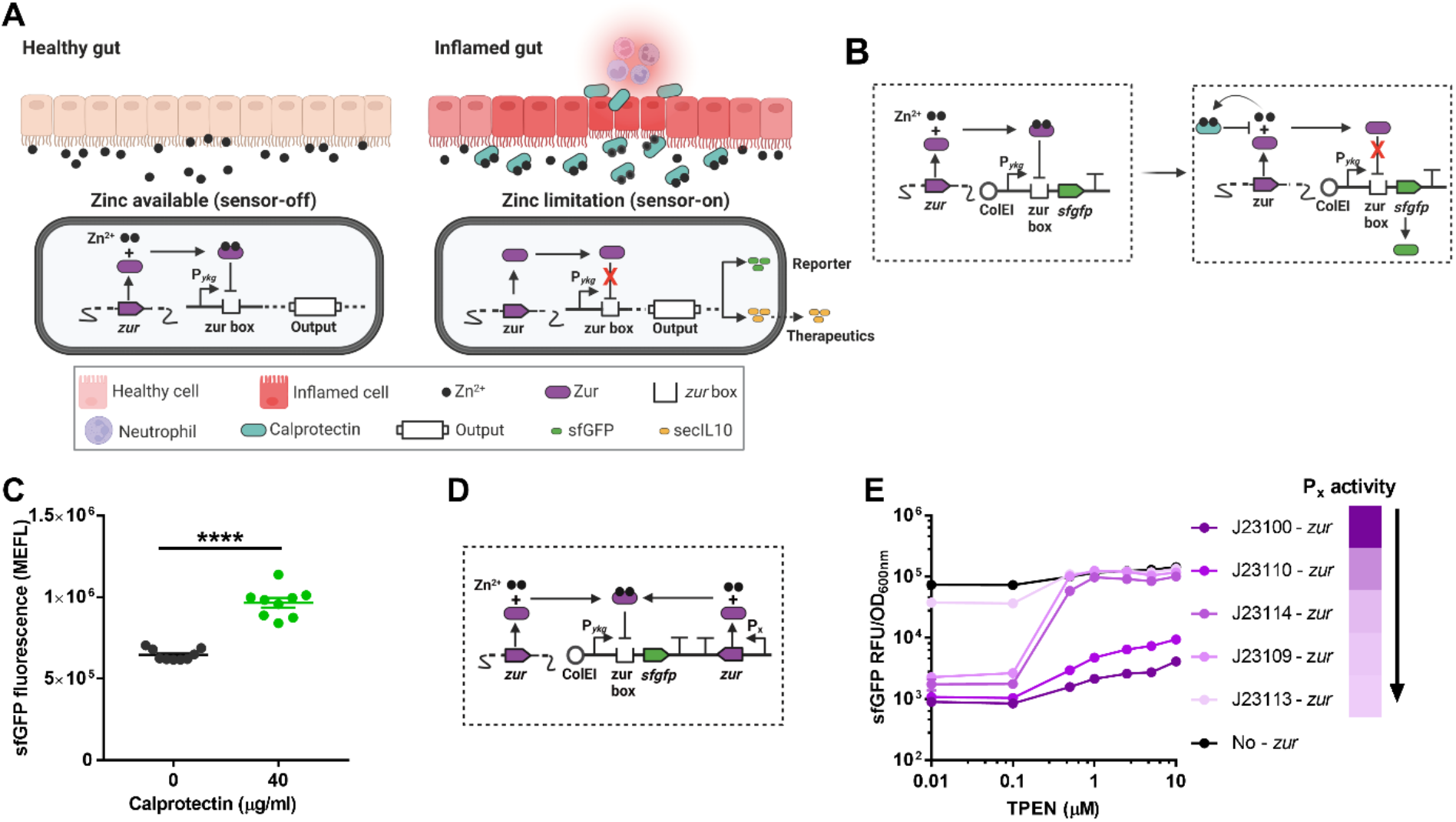
Identification and optimization of the *ykg* promoter construct to sense zinc limitation. **A:** Schematic diagram of engineered calprotectin-responsive biosensors for inflammation detection and therapeutic delivery. **B**: Schematic diagram of the biosensor with plasmid pBSI1 (PRB5000) responding to zinc limitation due to calprotectin. Left panel: zinc replete condition. Right panel: zinc depleted due to binding by calprotectin. **C**: Activation of biosensor BSI0 with 40 µg/ml calprotectin in M9 media. Data are mean ± SEM, n = 9 biological independent repeats. Statistical analysis was performed using an unpaired two-tailed t test; \*\*\*\**P* < 0.0001. **D**: Schematic diagram for testing promoters to identify the amount of Zur required for optimal dynamic range of the biosensor. P_x_ indicates where promoters of different strengths were integrated to alter the levels of *zur* in the cell. **E:** Optimization of the biosensor by altering the level of *zur* expression. Promoters that were tested are indicated on the right with the strength of the promoter indicated by the color gradient. The darkest color indicates the strongest promoter with decreasing strength moving down. Each construct was incubated in M9 medium supplemented with different amounts of TPEN (x-axis) and fluorescence was measured and normalized to the density of the culture to account for any slight differences in cell numbers (y-axis).

## Results

### Identification of a calprotectin sensitive promoter in *E. coli* Nissle

Bacteria use an evolutionarily conserved mechanism to regulate gene expression in response to zinc limitation that is dependent on the transcription factor Zur, which represses transcription in the presence of zinc-replete concentrations and is inactive when intracellular zinc levels fall below a critical threshold (*19, 20*). In *E. coli*, Zur controls the expression of two ribosomal protein genes, *ykgM* and *ykgO*, that encode homologs of the ribosomal proteins L31 and L36, which require zinc for their function in the ribosome. These alternative ribosomal proteins do not require zinc for function, are not expressed under normal conditions, and are highly expressed upon zinc depletion to allow ribosome function when zinc concentrations are low (*19*). These properties of the *ykgMO* operon made it an attractive candidate to engineer for an inflammation biosensor.

We cloned the *ykgMO* promoter (designated P_*ykg*_) of EcN upstream of the *sfgfp* gene to monitor expression in the presence of calprotectin (Fig. 1B). We incubated the resulting strain PRB5000 (EcN/pBSI1) in the presence of 40 µg/ml of calprotectin and observed a significant increase in *sfgfp* expression compared to untreated cells (Fig. 1C). To confirm this induction was due to zinc limitation, we also showed that the metal chelator Tetrakis-(2-pyridylmethyl) ethylenediamine (TPEN) induced PRB5000 induction (fig. S1A) and that the addition of zinc, but not iron, can restore repression of the promoter in the presence of calprotectin (fig. S1B).

### Optimization of P_*ykg*_ to sense zinc limitation

To increase the dynamic range of the P_*ykg*_ promoter, we postulated that the high basal level of expression was due to having P_*ykg*_ on a multicopy plasmid while *zur* was present in a single copy on the chromosome. To address this problem, we constructed a series of plasmids with *zur* under the control of a range of promoters with different strengths on the same plasmid as the P_*ykg*_-*sfgfp* construct (Fig. 1D). Two promoters, J23114 and J23109 were able to decrease basal expression from P_*ykg*_ ∼100-fold while retaining full induction by zinc limitation (Fig. 1E and fig. S1C). The strongest promoters tested successfully repressed P_*ykg*_ but did not allow full activation under zinc limitation while the weakest promoter had little effect on the basal level of transcription. We moved forward with J23109-*zur* and designated this construct pBSI2 and the strain BSI (BioSensor of Inflammation).

While calprotectin is a well-accepted biomarker for Ulcerative Colitis, there remains no generally accepted clinical biomarker for detecting inflammation non-invasively in Crohn’s disease patients. Calprotectin likely fails to identify inflammation accurately in Crohn’s patients due to the fact inflammation is patchy and often occurs in the small intestine, resulting in the loss of the calprotectin signal in feces (*21*). To create a bacterial biosensor that can both sense and remember encountering inflammation, we constructed a two-plasmid system in which the first plasmid is identical to pBSI1 but the expression of the gene integrase 8 (*int8*) is being driven by P_*ykg*_ (pBSIM1) (Fig. 2A). On the second plasmid, the *sfgfp* gene is placed in the opposite orientation of the strong promoter J23119 and flanked by *attB* and *attP* that are sites recognized by integrase 8 (pBSIM2). Expression of integrase 8 will flip the orientation of the *sfgfp* gene and allow for the constitutive expression of *sfgfp*. The EcN strain carrying both plasmids is referred to as BSIM (BioSensor of Inflammation with Memory).

**Fig. 2.**
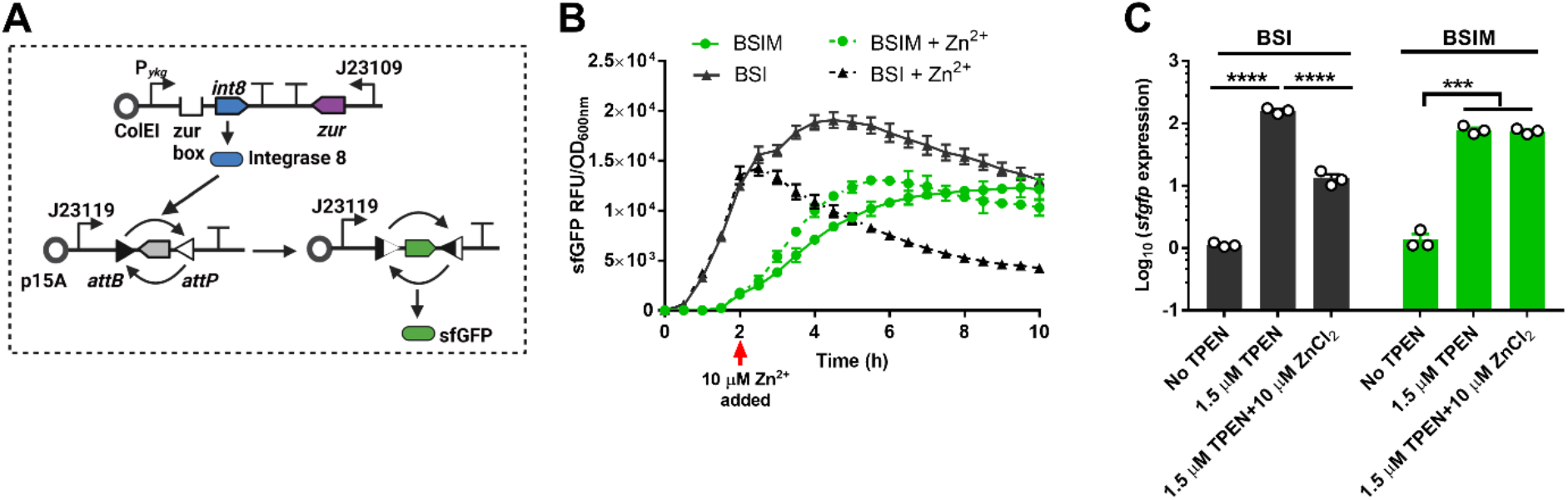
Genetic memory circuit biosensor senses and records zinc limitation. **A:** Schematic diagram of memory circuit biosensor BSIM. A two-plasmid system was created to allow for the biosensor to record that inflammation was detected. The first plasmid indicated in the top of the figure is the same as pBSI1 with the exception of the gene *int8* being placed under the control of P_*ykg*_. Expression of Integrase 8 leads to the inversion of the *sfgfp* gene in the second plasmid that allows for constitutive *sfgfp* expression. **B:** Stable expression of *sfgfp* in BSIM in Zn^2+^ replete conditions. BSI and BSIM were incubated in M9 medium with 1.5 µM TPEN to deplete zinc and induce biosensor activation. After 2 hours, 10 µM of zinc was added to the indicated cultures and fluorescence was measured and normalized. Data are mean ± SEM, n = 3 biological independent repeats. **C:** Expression of *sfgfp* in BSI and BSIM after Zn2+ addition. BSI and BSIM were incubated in M9 medium with 1.5 µM TPEN to deplete zinc and induce biosensor activation. After 2 hours, 10 µM of zinc was added. Total RNA from BSI and BSIM cultures was isolated four hours after zinc addition. Expression levels of *sfgfp* were measured by qRT-PCR. Data are mean ± SEM, n = 3 biological independent repeats. Statistical analysis was performed using ANOVA Tukey test; \*\*\**p* < 0.001.

To test the dynamics of induction of BSIM, we compared the expression of *sfgfp* in BSI1 and BSIM after treatment with TPEN. BSI immediately showed induction of sfGFP that steadily increased over the course of the experiment (Fig. 2B). For BSIM, expression of sfGFP is first observed two hours after TPEN addition, likely due to the time needed to express Int8 and reverse the orientation of *sfgfp* within the population. To demonstrate the memory function of BSIM, we added 10 µM of zinc to the cultures after two hours of induction. While BSI showed an immediate decrease in activity after zinc addition, BSIM continued to express sfGFP stably throughout the rest of the culture incubation and showed no difference in protein or RNA expression levels with or without the addition of zinc (Fig. 2B and 2C). We confirmed the biosensor was irreversibly activated by taking a subculture of the induced culture and incubating it in zinc replete conditions for multiple generations, observing constitutive expression of sfGFP (fig. S2).

### Detection of intestinal inflammation by BSIM

To demonstrate the utility of BSIM in accurately measuring inflammation *in vivo*, we tested the system in two independent mouse models of intestinal inflammation (Fig. 3). We treated mice with 1-3% dextran sulfate sodium (DSS) for six days to induce inflammation and then orally delivered BSIM (Fig. 3A). After four hours, colon and cecal contents were tested by qPCR to identify the change in the flipped orientation of the *sfgfp* gene, indicating biosensor activation. Increasing concentrations of DSS resulted in increased levels of calprotectin (Fig. 3B) and 3% DSS resulted in weight loss in the animals (fig. S3A). BSIM was significantly activated in the mice treated with 2 or 3% DSS (Fig. 3C) and the degree of activation was strongly correlated with calprotectin levels in the intestine (Fig. 3D). We also isolated several hundred colonies from each treatment and showed that a significantly higher proportion of isolated colonies had *sfgfp* activated in 3% DSS treated animals compared to healthy controls (fig. S3B). These results demonstrate that BSIM can accurately distinguish between healthy and inflamed mice in a calprotectin dependent manner.

**Fig. 3.**
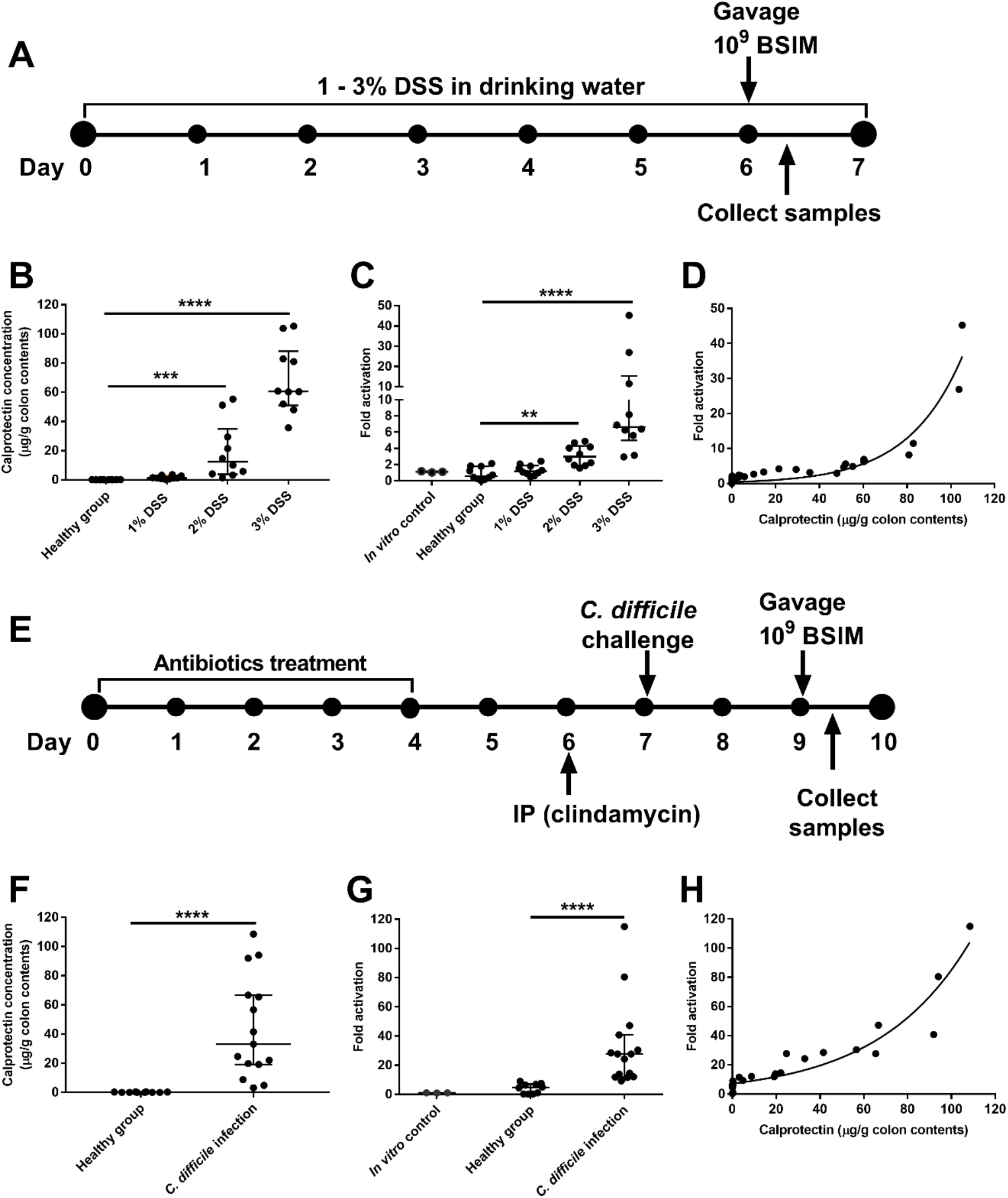
Detection of intestinal inflammation by BSIM. **A:** Experimental design for testing BSIM in DSS-induced gut inflammation mouse model. 6-week-old C57BL/6 mice (5 female and 5 male mice in each group) were given water with or without 1, 2 or 3% DSS for 6 days before oral gavage with BSIM. After 4 h, samples were collected from the mice and processed for the next analysis. **B:** Concentration of calprotectin in mouse colon contents. Colon contents at the end of the experiment were isolated and tested for calprotectin levels by ELISA. Data are median ± interquartile range, n = 10; Individual dots represent individual mice. Statistical analysis was performed using ANOVA Kruskal-Wallis test; \*\*\**p* < 0.001, \*\*\*\**P* < 0.0001. **C:** Fold activation of BSIM in mouse colon contents. Total RNA was isolated from colon contents and the orientation of the *sfgfp* gene was detected by qRT-PCR. Fold increase in the flipped orientation of *sfgfp* compared to the *in vitro* control is presented. Data are median ± interquartile range; for *in vitro* control n = 3 biological independent repeats, for *in vivo* experiments n = 10, individual dots represent individual mice. Statistical analysis was performed using ANOVA Kruskal-Wallis test; \*\**p* < 0.01, \*\*\*\**P* < 0.0001. **D:** Correlation analysis of biosensor BSIM activation and colon calprotectin concentration. n = 40; individual dots represent individual mice. Correction analysis was performed by computing Pearson correlation coefficients with 95% confidence interval and two tailed *p* value analysis. (95% CI = 0.616 to 0.877, *r* = 0.778, *p* < 0.0001). **E:** Experimental design for testing BSIM in *C. difficile* induced gut inflammation mouse model. Briefly, 6-week-old C57BL/6 mice (for the healthy group n = 10 with 5 female and 5 male mice, for the *C. difficile* infection group n = 20 with 10 female and 10 male mice) were treated with five mixed antibiotics cocktail for four days, following intraperitoneal injection of one dose of clindamycin and oral gavage of 10^5^ spores of strain *C. difficile* R20291. Two days post infection, mice were gavaged with 10^9^ CFU of BSIM. After 4 h, colon samples were collected and processed for analysis. **F:** Concentration of calprotectin in the mice colon contents. Colon contents at the end of the experiment were isolated and tested for calprotectin levels by ELISA. Data are median ± interquartile range, for the healthy group n = 10, for the *C. difficile* infection group n = 15 (five mice were died or difficult to get samples); Individual dots represent individual mice. Statistical analysis was performed using Mann-Whitney test; \*\*\*\**P* < 0.0001. **G:** Fold activation of BSIM in mouse colon contents. Total RNA was isolated from colon contents and the orientation of the *sfgfp* gene was detected by qRT-PCR. Fold increase in the flipped orientation of *sfgfp* compared to the *in vitro* control is presented. Data are median ± interquartile range, healthy group n = 10, *C. difficile* infection group n = 15; Individual dots represent individual mice. Statistical analysis was performed using Mann-Whitney test; \*\*\*\**P* < 0.0001. **H:** Correlation of biosensor BSIM activation and colon calprotectin concentration. n = 25; Individual dots represent individual mice. Correction analysis was performed by computing Pearson correlation coefficients with 95% confidence interval and two tailed *p* value analysis. (95% CI = 0.798 to 0.959, *r* = 0.907, *p* < 0.0001).

To further demonstrate the ability of BSIM to sense inflammation in the intestine, we used the *Clostridium difficile* infection model that has been shown to induce a high level of neutrophil invasion and calprotectin release (*22*). Mice were treated with antibiotics and infected with R20291, a ribotype 027 strain of *C. difficile* that causes moderate to severe disease (*23, 24*). After two days of *C. difficile* infection the mice were gavaged with BSIM and biosensor activation was monitored as described above (Fig. 3E). We found that *C. difficile* infection caused a significant increase in calprotectin (Fig. 3F) and activation of the BSIM occurred in mice infected with *C. difficile* (30-fold increase) (Fig. 3G). As observed in the DSS model, the level of activation (Fig. 3H) had a strong correlation with the level of calprotectin detected in the colon. These results indicate the BSIM system is accurately detecting levels of inflammation in the context of an inflamed gut in two independent models.

### Biosensor coupling with a therapeutic to sense and respond to intestinal inflammation

To ask if our bacterial biosensors could be re-designed to sense inflammation and deliver an anti-inflammatory therapeutic in response, we replaced the *sfgfp* gene in pBSI2 with the human IL-10 gene fused to the YebF protein (referred to hereafter as secIL10) resulting in therapeutic sensor PRB5003 (EcN/PBSIDZ3) (Fig. 4A). YebF has been shown to facilitate the secretion of proteins in *E. coli* (*18*). Induction of P_*ykg*_ with TPEN yielded ∼40 ng/ml/OD of secIL10 into the extracellular media after four hours of induction (Fig. 4B). IL-10 fused to another secretion signal yielded only ∼40 pg/ml/OD of IL10 (Fig. S4C). We next confirmed that secIL10 has functional IL10 activity using a cell line (Human & Murine IL-10 reporter cells) that couples IL10 receptor activation to a colorimetric reporter output. In comparison to recombinant IL10, secIL10 displayed a similar level of IL10 activity indicating the fusion is functional (Fig. 4C).

**Fig. 4.**
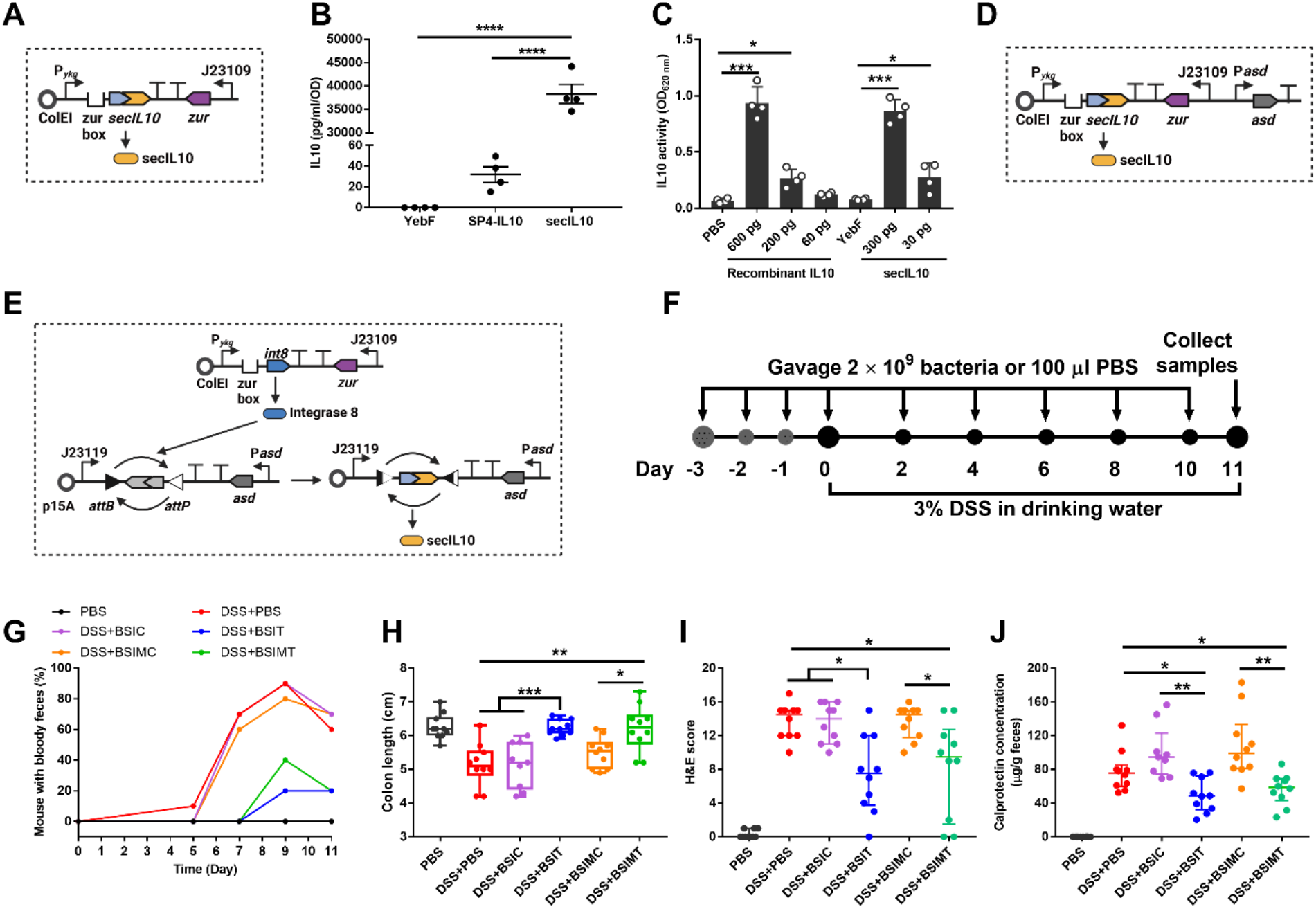
Biosensor coupling with a functional human IL10 senses and responds to intestinal inflammation. **A:** Diagram of therapeutic sensor plasmid pBSIDZ3. The *yebF* gene from EcN was fused into the N-terminal of *IL10* (referred to as secIL10) and placed under the control of P_*ykg*_. **B:** Fusion of YebF to IL10 results in high level secretion of secIL10. Biosensor PRB5001 (EcN/PBSIDZ1) and PRB5003 (EcN/PBSIDZ3) were cultured to OD_600_ of 0.5 - 0.6 and then induced with 20 µM TPEN for 4 h in LB media. Following, the supernatants of cultures were collected and detected by IL10 ELISA kit. Control sensor PRB5002 (EcN/PBSIDZ2) that expresses YebF only was used as a control. Data are mean ± SEM, n = 4 biological independent repeats. Statistical analysis was performed using ANOVA Tukey test; \*\*\*\**p* < 0.0001. **C:** Testing the activity of secIL10 with HEK-Blue IL10 reporter cell line. Quantified secIL10 from BSIT supernatant was added to HEK-Blue IL-10 reporter cells line to activate reporter expression. Recombinant human IL10 was used as a positive control. Supernatant from PRB5002 (EcN/PBSIDZ2) with only YebF were used as a negative control. Data are mean ± SEM, n = 4 biological independent repeats; Statistical analysis was performed using ANOVA with Tukey’s multiple-comparisons test; \**p* < 0.05,\*\*\**p* < 0.001. **D-E:** Diagram of secIL10 stable expression sensor BSIT (Δ*asd*/pBSIT) and BSIMT (Δ*asd* with pBSIM1 and pBSIMT plasmids). The *asd* gene with its own promoter from EcN was cloned into pBSIT and pBSIMT plasmid, respectively. **F:** Experimental design of therapeutic biosensors applied to DSS-induced gut inflammation for the treatment of intestinal inflammation. 6-week-old C57BL/6 mice were orally gavaged 2×10^9^ therapeutic sensor bacteria or 100 µl PBS for three days, following mice were treated with or without 3% DSS for 11 days. 2×10^9^ therapeutic sensor bacteria or 100 µl PBS were gavaged every two days. On day 11, mice were euthanized and colon samples were collected. One group of mice was gavaged with PBS without DSS treatment as a control. **G:** Percentage of mice with bloody feces. Mice producing bloody feces were monitored and recorded every day for 11 days. Groups tested include DSS + BSIT, DSS + BSIMT, DSS + BSIC (control expressing only YebF), DSS + BSIMC (control expressing only YebF) and DSS + PBS (colitis control). Groups were compared to animals that received only gavage of PBS without DSS treatment. **H:** Colon length. Data are presented in a box & whiskers plot with min to max, show all points, n = 10; Individual dots represent individual mice. Statistical analysis was performed using ANOVA Kruskal-Wallis test; \**P* < 0.05, \*\**P* < 0.01, \*\*\**P* < 0.001. **I:** Histological score of colon tissue. Mouse distal colon tissue was collected, fixed, sliced, and stained with hematoxylin & eosin (H&E staining). A blinded histological scoring was performed. Six inflammation related items (inflammatory infiltrate, goblet cell loss, crypt hyperplasia, muscle thickening, submucosal inflammation, ulceration) were evaluated with each item from 0-3 score based on disease severity. Data are median ± interquartile range, n = 10; Individual dots represent individual mice. Statistical analysis was performed using ANOVA Kruskal-Wallis test; \**P* < 0.05. **J:** Concentration of calprotectin in mice feces. Mice fecal samples were collected on day 11 and tested for calprotectin concentration by ELISA. Data are median ± interquartile range, n = 10; Individual dots represent individual mice. Statistical analysis was performed using ANOVA Kruskal-Wallis test; \**P* < 0.05.

Because plasmids will not be under selection while in the mouse intestine, we engineered into the system the expression of the *asd* gene, which encodes the enzyme aspartate semialdehye dehydrogenase, that is required for lysine, threonine, and methionine biosynthesis. Disruption of *asd* in the *E. coli* Nissle chromosome (?*asd*) makes growth of the strain dependent on retention of the plasmid with *asd* gene (Fig. 4D). We confirmed that addition of *asd* to the pBSIDZ3 (pBSIT) resulted in significant retention of the plasmid in the absence of selection (fig. S5A), as previously described (*7*). We refer to this system as BSIT (BioSensor of Inflammation with Therapeutic). We made the same modifications to the BSIM plasmid constructs resulting in BSIMT (Fig. 4E) (BioSensor of Inflammation with Memory and Therapeutic) and validated a similar level of secIL10 induction (fig. S5B).

To test if BSIT and BSIMT could sense, respond, and alleviate intestinal inflammation, we treated mice with 3% DSS with BSIT, BSIMT, and controls of *E. coli* Nissle only expressing YebF (designated BSIC and BSIMC). We gavaged animals with each strain for three days prior to starting DSS treatment and then provided the organisms every other day during 3% DSS treatment (Fig. 4F). Animals receiving only PBS vehicle were used as an additional negative control. We found that BSIT and BSIMT were able to significantly improve several parameters associated with DSS colitis including reduction in mice with bloody feces (Fig. 4G), improving colon length (Fig. 4H), and improved histological scores compared to the PBS, BSIC and BSIMC treated controls (Fig. 4I). We also found that BSIT and BSIMT treatment resulted in reduced levels of calprotectin in the colon, consistent with reduced intestinal inflammation (Fig. 4J). These data show that both systems can sense intestinal inflammation and resolve the inflammation by production of an anti-inflammatory protein.

## Discussion

An important finding of this work is that the P_*ykg*_ promoter is naturally tuned to respond to the levels of zinc limitation that occur during neutrophil infiltration and inflammation in the gut. We posited this would be the case as *E. coli* has co-evolved with humans for millennia and the system was optimized to respond to nutritional immunity. We expect this occurs because zinc levels need to reach extremely low levels to render Zur inactive, and this likely occurs only at sites of inflammation. The strong correlation between colonic calprotectin levels and biosensor activation in colitis indicates that rather than being an all or nothing response there is the ability to sense different levels of inflammation.

Our memory circuit biosensor is a prototype for detecting the patchy inflammation that occurs in the small intestine in Crohn’s disease. Currently there is no good biomarker for detecting inflammatory signals in feces in Crohn’s disease and the memory biosensor BSIM may be able to fill this need. One hurdle that needs to be overcome is the ability to measure the level of activation without the need for the handling of feces. Coupling the output of BSIM with the ability to produce a color change in the stool or another metabolic readout that can be monitored in the toilet water is needed. The role of Zur in the regulation of ribosomal proteins involved in zinc dependent regulation is evolutionarily conserved and is found in many bacteria (*26, 27*). We expect these promoters in other organisms will be of similar value in developing additional inflammation biosensors for the gut and other sites of the body.

The coupling of the sensing of inflammation to production of IL-10 demonstrated that rather than having constitutive expression of a therapeutic expressed from a synthetic organism that it is possible to express these proteins when disease is encountered. As many immunologic treatments for IBD and other inflammatory diseases are usually systemic and have unwanted side effects associated with body wide immunosuppression, local delivery is likely able to circumvent some of these side effects. Our sense and respond system provides further control of expression of therapeutics to only occur when disease is encountered, further reducing the chances of side effects associated with systemic therapeutic delivery.

## Acknowledgements

We thank Dr. Annie Goodwin and Dr. Shuai Qian for technical assistance in developing and testing the microbial biosensor and Dr. Michael Ittmann for blinded histological scoring. We thank Dr. Walter Chazin for providing functional recombinant calprotectin. We thank Dr. Vincent Young for critical review of the manuscript. We thank members of the Britton lab for valuable feedback on this work.

## Author Contributions

Conceptualization: RAB

Methodology: DA, JDG, JP, EM, JJT, RAB

Investigation: DZ, JDG

Supervision: JJT, RAB

Writing – Original draft: DZ, RAB

Writing – Review and editing: DZ, JDG, JJT, RAB

## Data and materials availability

All data are available in the main text or the supplementary materials.

## Supplementary Materials

Materials and Methods

Figs. S1 to S5

Tables S1 to S2

References (*6, 28*)

## References

1. A. Cubillos-Ruiz et al., Engineering living therapeutics with synthetic biology. Nat Rev Drug Discov 20, 941–960 (2021).

2. M. R. Charbonneau, V. M. Isabella, N. Li, C. B. Kurtz, Developing a new class of engineered live bacterial therapeutics to treat human diseases. Nat Commun 11, 1738 (2020).

3. B. P. Landry, J. J. Tabor, Engineering Diagnostic and Therapeutic Gut Bacteria. Microbiol Spectr 5, (2017).

4. D. T. Riglar, P. A. Silver, Engineering bacteria for diagnostic and therapeutic applications. Nat Rev Microbiol 16, 214–225 (2018).

5. N. Aggarwal, A. M. E. Breedon, C. M. Davis, I. Y. Hwang, M. W. Chang, Engineering probiotics for therapeutic applications: recent examples and translational outlook. Curr Opin Biotechnol 65, 171–179 (2020).

6. K. N. Daeffler et al., Engineering bacterial thiosulfate and tetrathionate sensors for detecting gut inflammation. Mol Syst Biol 13, 923 (2017).

7. T. Chien et al., Enhancing the tropism of bacteria via genetically programmed biosensors. Nat Biomed Eng 6, 94–104 (2022).

8. L. Liu et al., An Electrochemical Biosensor with Dual Signal Outputs: Toward Simultaneous Quantification of pH and O(2) in the Brain upon Ischemia and in a Tumor during Cancer Starvation Therapy. Angew Chem Int Ed Engl 56, 10471–10475 (2017).

9. N. E. Walsham, R. A. Sherwood, Fecal calprotectin in inflammatory bowel disease. Clin Exp Gastroenterol 9, 21–29 (2016).

10. A. Ricciuto, A. M. Griffiths, Clinical value of fecal calprotectin. Crit Rev Clin Lab Sci 56, 307–320 (2019).

11. T. Vogl, N. Leukert, K. Barczyk, K. Strupat, J. Roth, Biophysical characterization of S100A8 and S100A9 in the absence and presence of bivalent cations. Biochim Biophys Acta 1763, 1298–1306 (2006).

12. M. Fagerhol, K. Andersson, C. Naess-Andresen, P. Brandtzaeg, I. J. B. R. Dale, Fla: CRC Press, Inc, Calprotectin (the L1 leukocyte protein) In: Smith VL, Dedman JR, editors. Stimulus response coupling: the role of intracellular calcium-binding proteins. 187–210 (1990).

13. M. B. Brophy, J. A. Hayden, E. M. Nolan, Calcium ion gradients modulate the zinc affinity and antibacterial activity of human calprotectin. J Am Chem Soc 134, 18089–18100 (2012).

14. T. G. Nakashige, B. Zhang, C. Krebs, E. M. J. N. c. b. Nolan, Human calprotectin is an iron-sequestering host-defense protein. 11, 765–771 (2015).

15. T. G. Nakashige, E. M. Zygiel, C. L. Drennan, E. M. Nolan, Nickel Sequestration by the Host-Defense Protein Human Calprotectin. J Am Chem Soc 139, 8828–8836 (2017).

16. M. Schultz, Clinical use of E. coli Nissle 1917 in inflammatory bowel disease. Inflamm Bowel Dis 14, 1012–1018 (2008).

17. J. P. Lynch, L. Goers, C. F. Lesser, Emerging strategies for engineering Escherichia coli Nissle 1917-based therapeutics. Trends Pharmacol Sci 43, 772–786 (2022).

18. G. Zhang, S. Brokx, J. H. Weiner, Extracellular accumulation of recombinant proteins fused to the carrier protein YebF in Escherichia coli. Nat Biotechnol 24, 100–104 (2006).

19. S. E. Gabriel, J. D. Helmann, Contributions of Zur-controlled ribosomal proteins to growth under zinc starvation conditions. J Bacteriol 191, 6116–6122 (2009).

20. J. Z. Liu et al., Zinc sequestration by the neutrophil protein calprotectin enhances Salmonella growth in the inflamed gut. Cell Host Microbe 11, 227–239 (2012).

21. C. McDowell, M. Haseeb, Inflammatory bowel disease (IBD). (2017).

22. S. Jose, R. Madan, Neutrophil-mediated inflammation in the pathogenesis of Clostridium difficile infections. Anaerobe 41, 85–90 (2016).

23. A. M. Buckley, J. Spencer, D. Candlish, J. J. Irvine, G. R. Douce, Infection of hamsters with the UK Clostridium difficile ribotype 027 outbreak strain R20291. J Med Microbiol 60, 1174–1180 (2011).

24. D. Zhu, J. Bullock, Y. He, X. Sun, Cwp22, a novel peptidoglycan cross-linking enzyme, plays pleiotropic roles in Clostridioides difficile. Environ Microbiol 21, 3076–3090 (2019).

25. S. Han et al., Novel signal peptides improve the secretion of recombinant Staphylococcus aureus Alpha toxin(H35L) in Escherichia coli. AMB Express 7, 93 (2017).

26. E. M. Panina, A. A. Mironov, M. S. Gelfand, Comparative genomics of bacterial zinc regulons: enhanced ion transport, pathogenesis, and rearrangement of ribosomal proteins. Proc Natl Acad Sci U S A 100, 9912–9917 (2003).

27. D. Kandari, H. Joshi, R. Bhatnagar, Zur: Zinc-Sensing Transcriptional Regulator in a Diverse Set of Bacterial Species. Pathogens 10, (2021).

28. Y. Jiang et al., Multigene editing in the Escherichia coli genome via the CRISPR-Cas9 system. Appl Environ Microbiol 81, 2506–2514 (2015).

