## Supplementary material for "Microbial biosensor for sensing and treatment of intestinal inflammation"

Duolong Zhu *et al.*

**The PDF file includes:**

Materials and Methods

Figs. S1 to S5

Tables S1 to S2

References

### Materials and Methods

#### Bacterial strains and media

All strains used are listed in Table S1. *Escherichia coli* Nissle 1917 (referred hereafter as EcN) was grown aerobically at 37 °C using either Luria-Bertani (LB) broth (1% tryptone, 0.5% yeast extract, 1% NaCl) or the minimal defined media M9 (200 ml 5×M9 salts+0.4% glucose+2 mM $MgSO_4$+0.1 mM $CaCl_2$+0.2% casamino acids for 1 liter M9 media). *Escherichia coli* DH5α was used as a cloning host. Chloramphenicol (15 µg/ml) and ampicillin (100 µg/ml) were used for *E. coli* selection when needed. *Clostridioides difficile* R20291 was cultured in BHIS media (brain heart infusion broth supplemented with 0.5% yeast extract and 0.1% L-cysteine, and 1.5% agar for agar plates) at 37 °C in an anaerobic chamber (90% $N_2$, 5% $H_2$, 5% $CO_2$). For spores preparation, *C. difficile* strains were cultured in Clospore media and purified as described earlier (*27*).

#### Biosensor plasmids construction and optimization

The *ykgMO* (paralog for L31/L36 ribosomal accessory protein) promoter was cloned directly upstream of a super fold green fluorescent protein cassette (*sfgfp*) in pColE1 plasmid via the Gibson Reaction (New England Biosciences, Ipswich, Massachusetts). The resulted plasmid and biosensor strain were named as pBSI1 (pColE1-P*ykg*- *sfgfp*) and PRB5000, respectively.

To reduce the background of primary P*ykg* biosensor, zinc binding transcription factor *zur* gene was inserted into PRB5000 under the control of different constitutive expression promoters. Among them, five different constitutive promoters J23100, J23110, J23114, J23109, and J23113 from Registry of Standard Biological Parts (http://parts.igem.org/Promoters/Catalog/Anderson) were selected and tested. The sensitivity and expression strength of new biosensor constructs was analyzed through fluorescence detection.

Because we expect that zinc sequestration to levels that will activate the biosensor will only occur at sites of active inflammation with neutrophil infiltration, we posited that the ideal biosensor would need to become permanently activated upon sensing inflammation for future readout in the stool. We developed a memory switch biosensor containing a two-plasmid system using phage integrase 8 being driven by the $P_{ykg}$ promoter regulated by Zur on one plasmid (pBSIM1) and a *sfgfp* gene on the second plasmid in the opposite orientation of the strong promoter J23119 (denoted pBSIM2). The *sfgfp* gene is flanked by *att* sites that are recognized by the integrase and when expressed will flip the orientation of the *sfgfp* gene to allow expression from promoter J23119. All Gibson oligos were made using IDT, and all amplicons were synthesized with Phusion polymerase (NEBiosciences, Ipswich, Massachusetts). A listing of all constructed plasmids and primers used in this study can be found in Table S1 and Table S2.

#### Flow cytometry and data analysis

EcN biosensor constructs were grown to early log phase (~0.1 $OD_{600}$) in M9 media and were back diluted to 0.01 $OD_{600}$ and then grown for ~4 hours aerobically, until $OD_{600}$ reached ~0.15. At this point, cells were kept on ice until analysis on a Becton Dickinson FACScan flow cytometer. Flow cytometry runs were performed in 96 well plates. Briefly, 10 - 40 µl of cells were added to 1 ml of PBS sheath fluid and run through the flow cytometer for a total of 10000 events. Cells were thresholded on forward/side scatter, and .fsc files were analyzed with the FlowCal software, developed by the Jeff Tabor Lab. In short, FlowCal identifies the densest region of cells on the associated scatterplot and analyzes the fluorescent output of 30% of the cells in this region so as to evaluate a homogenous dataset and remove possible outliers (*6*). This results in a geometric mean of total fluorescent output. In this case, *sfgfp* output is reported as molecules of equivalent fluorophores (MEF) and was evaluated on the FL1 channel. More information on FlowCal can be found at: http://taborlab.github.io/FlowCal/.

**Calprotectin and TPEN Induction and metal complementation**

Final concentrations of 40 µg/ml recombinant human calprotectin and 1.5, 3, or 30 µM TPEN (*N*,*N*,*N'*,*N'*-tetrakis(2-pyridinylmethyl)-1,2-ethanediamine) were used for the biosensor constructs induction test, respectively. The following mix was added to each well- 124 µl of recombinant human calprotectin in calprotectin buffer, 56 µL of M9 media, and 20 µl of cells ($10^4$). TPEN (Sigma Aldrich, St. Louis, MO), a synthetic metal chelator, induction assays were performed as calprotectin induction, with the following differences- 160 µL of M9 media, 20 µl of cells ($10^4$), and 20 µl of either 15, 30, or 300 µM TPEN was added to each well. TPEN was dissolved in absolute ethanol. The absolute ethanol was used as a negative control.

Mixes were similar to the calprotectin induction assays with the addition of zinc sulfate, manganese sulfate, or iron chloride. Concentrations of metals were added in excess of 1× and 10× the binding capacity of 40 µg/ml (1.5 µM) calprotectin. In total, 4 µM and 40 µM of zinc and iron were added, and 2 µM and 20 µM of manganese was added.

**Therapeutic biosensor construction**

To engineer human IL10 secretion strain, signal peptide NSP4 and facilitating carrier protein YebF were fused up to IL10 and assembled into plasmid pBSI2 which replaced *sfgfp*, respectively. The corresponding therapeutic strains were denoted as PRB5001 (with pBSIDZ1 plasmid) and PRB5003 (with pBSIDZ3 plasmid).

To achieve stable expression of secIL10, an essential gene *asd* that is required for lysine, threonine, and methionine biosynthesis in EcN was deleted by CRISPR-Cas9 (*28*) and complemented the *asd* gene in sensor plasmid pBSIT, resulting a therapeutic sensor BSIT. Meanwhile, the *sfgfp* gene in pBSIM2 was replaced with secIL10, and the *asd* gene was assembled into pBSIM2 to get the stable therapeutic biosensor with a memory circuit (BSIMT). Strains with only *yebF* expression plasmid (pBSIC and pBSIMC) were used as controls (BSIC and BSIMC). Supernatants of therapeutic biosensor cultures with TPEN induced were analyzed by IL10 ELISA kit (EAGLE Biosciences, Nashua, NH). The activity of secIL10 was analyzed by the Human&Murine IL-10 reporter cells (HEK-Blue™ IL10 Cells) that is engineered to respond to functional IL-10 according to the product instruction (InvivoGen, San Diego, CA).

### Animal experiments

For dextran sodium sulfate (DSS, MW = 40,000, Thermos Scientific) induced IBD mouse model, six-week-old C57BL/6 mice were procured from Baylor College of Medicine in Houston, Texas. Mice were transferred to an established protocol that was approved by the Baylor College of Medicine Institutional Animal Care and Use Committee (IACUC). Mice were treated with or without 1 - 3% (w/v) DSS in drinking water for seven days. On the six day, mice were gavaged with $10^9$ CFU of memory circuit biosensor. 4 - 6 hours after gavage, fecal pellets and colon contents from all mice were collected. The total DNA from stool and colon contents were isolated by E.Z.N.A Stool DNA kit (Omega) according to the instruction for biosensor activation efficiency analysis by qPCR.

To test biosensor in the *Clostridioides difficile* infection mouse model (CDI), six-week-old C57BL/6 mice were given an orally administered antibiotic cocktail (kanamycin 0.4 mg $ml^{-1}$, gentamicin 0.035 mg $ml^{-1}$, colistin 0.042 mg $ml^{-1}$, metronidazole 0.215 mg $ml^{-1}$, and vancomycin 0.045 mg $ml^{-1}$) in drinking water for 4 days. After 4 days of antibiotic treatment, all mice were given autoclaved water for 2 days, followed by one dose of clindamycin (10 mg $kg^{-1}$, intraperitoneal route) 24 h before the spores challenge (Day 0). After that, mice were orally gavaged with $10^{4-5}$ of spores or PBS as a control. 2 days after spores gavage, $10^9$ CFU of memory circuit biosensor were gavaged to all mice. 4-6 hours after biosensor gavage, fecal pellets and colon contents were collected for biosensor activation test.

To evaluate the therapeutic biosensor for inflammation amelioration *in vivo*, the IBD animal model was used. Six-week-old C57BL/6 mice were purchased. Before DSS treatment, mice were gavaged with PBS (control group and +DSS group) or therapeutic sensor constructs for 3 days. Following, the mice were treated with or without 3% (w/v) DSS in drinking water and orally gavaged with either 100 µl of PBS or $2\times10^9$ CFU of the therapeutic sensor every two days for 10 days. The mice weight and disease severity was monitored every day. On the day 11, mice were euthanized and fecal pellets, colon, and colon contents were collected.

### Supplementary figures

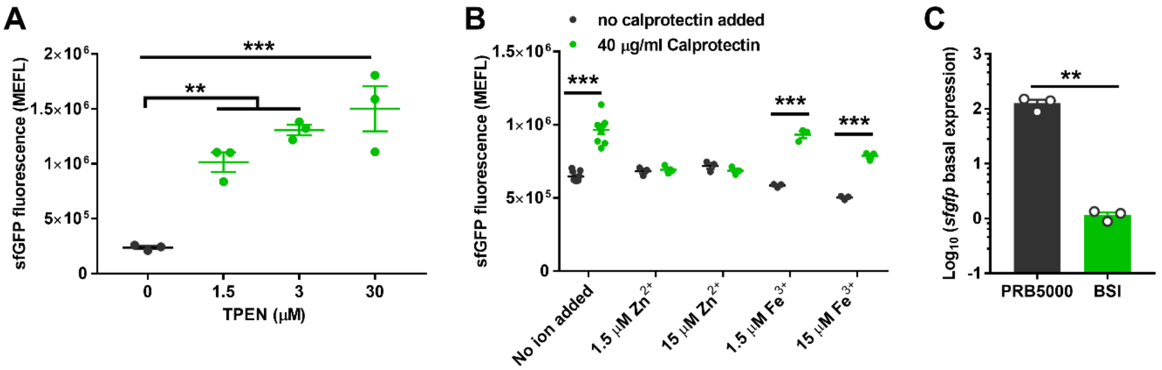


**fig. S1. Identification and optimization of the *ykg* promoter construct to sense zinc limitation.** **A:** Fluorescence of primary biosensor PRB5000 (EcN/pBSI1) induced by different concentrations of TPEN. 1.5 µM (equivalent metal binding capacity of 40 µg/ml calprotectin), 3 µM, or 30 µM TPEN was used to activate biosensor BSI0 in M9 media. Data are mean ± SEM, n = 3 biological independent repeats. Statistical analysis was performed using ANOVA Tukey test; $^{**}p < 0.01$, $^{***}p < 0.001$.

**B:** Zinc addition turns off sensor PRB5000 activation. Equimolar (1.5 µM) or 10 × fexcess (15 µM) of $Zn^{2+}$ and $Fe^{2+}$ were added to BSI0 co-cultured with 40 µg/ml calprotectin in M9 media. Data are mean ± SEM, for no ion added test n = 9 biological independent repeats, for metals added test n = 3 biological independent repeats. Statistical analysis was performed using ANOVA Tukey test; $^{***}p < 0.001$.

**C**: Basal expression of *sfgfp* in biosensor PRB5000 and BSI (EcN/pBSI2) detected by RT-qPCR. PRB5000 and BSI biosensors were cultured for 6 h, then total RNA was isolated for *sfgfp* gene transcription analysis. Data are mean ± SEM, n = 3 biological independent repeats. Statistical analysis was performed using an unpaired two-tailed t test; $^{**}p < 0.01$.

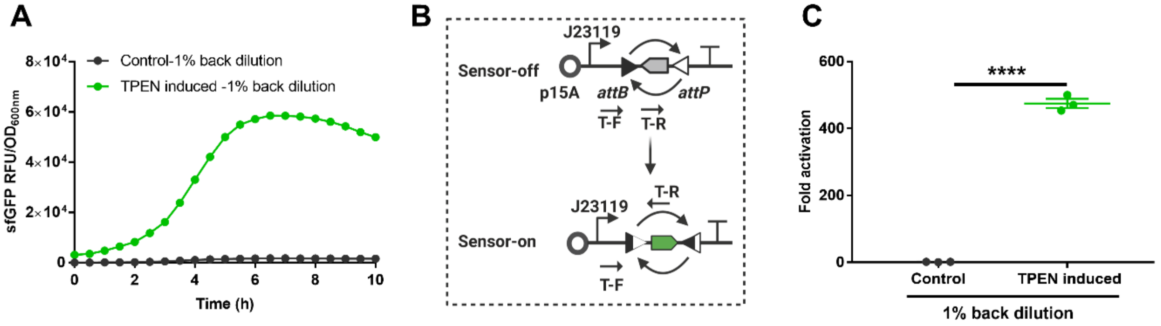


**fig. S2 Stable expression of *sfgfp* in memory circuit biosensor BSIM.**

**A**: The BSIM biosensor was induced with 1.5 μM TPEN for 4 h, then 1% of cultures were subculutred into fresh M9 media without TPEN. Fluorescence was monitored every 30 min for 10 hours. BISM without TPEN induction was used as a control. Data showed here was from the fifth passage. Data are mean ± SEM, n = 3 biological independent repeats.

**B**: Schematic diagram of BSIM sensor off and sensor on. Primers T-F/T-R were used for the flipped *sfgfp* gene detection. When *sfgfp* is on the reverse orientation (sensor off status), T-F/T-R primers cannot amplify the PCR bands.

**C**: Fold change of flipped *sfgfp* (fold activation) in 1% back dilution cultures at 8 h growth. Plasmids from 1% back dilution cultures were isolated and used for qPCR test with primers T-F/T-R. The ampicillin resistance gene (*amp*) on the sensor plasmid pBSIM2 was used as a control gene. Data are mean ± SEM, n = 3 biological independent repeats. Statistical analysis was performed using an unpaired two-tailed t test; *****P* < 0.0001.

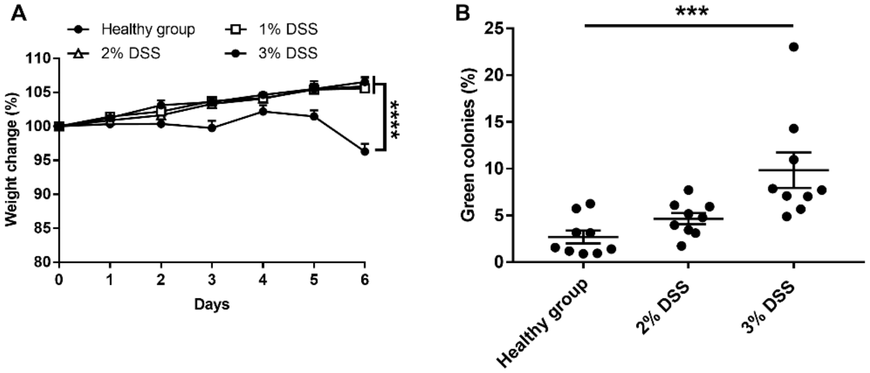


**fig. S3 Detection of intestinal inflammation by BSIM.**

**A:** Weight change of mice with treated with 1-3% DSS. Data are mean ± SEM, n = 10 for each group; Individual dots represent individual mice. Statistical analysis was performed using a paired two-tailed t test; ****$P < 0.0001$.

**B:** Percentage of bacterial colonies showing BSIM biosensor activation (green colonies) in the mouse colon detected by plating. Colon contents from healthy, 2%, and 3% DSS treated groups were serially diluted in PBS and plated on LB agar plates with 15 µg/ml chloramphenicol and 100 µg/ml ampicillin to select cells harboring BSIM. Colonies were analyzed by fluorescence microscopy and the percentage of green colonies per total colonies is reported. Data are mean ± SEM**,** n = 9; Individual dots represent individual mice. Statistical analysis was performed using ANOVA Kruskal-Wallis test; ***$P < 0.001$.

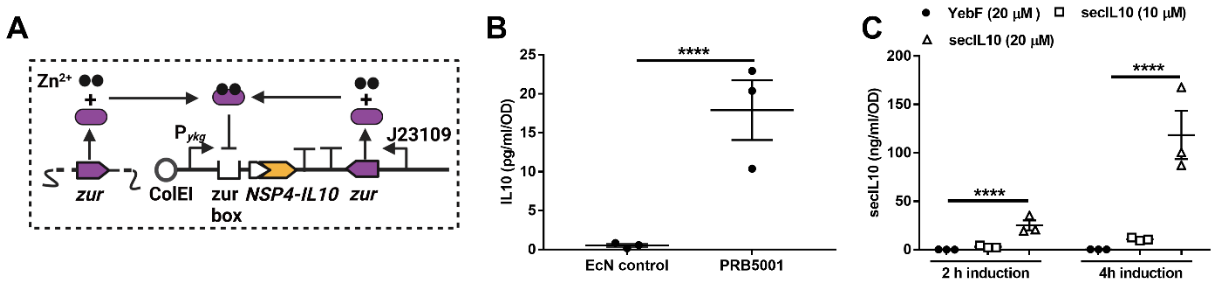


**fig. S4 Therapeutic sensor coupling with a human IL10.**

**A:** Diagram of therapeutic sensor plasmid pBSIDZ1. The human IL10 gene was fused with an *E. coli* secretion signal peptide NSP4 (NSP4-IL10).
**B:** Secretion of IL10 in biosensor PRB5001 (EcN/pBSIDZ1). Biosensor PRB5001 was cultured to $OD_{600}$ of 0.5 - 0.6 and then induced with 20 µM TPEN for 4 h in LB media. Following, the supernatants of cultures were collected and detected by IL10 ELISA kit. EcN supernatants was used as a control. Data are mean ± SEM, n = 3 biological independent repeats. Statistical analysis was performed using an unpaired two-tailed t test; $^{****}P < 0.0001$.
**C:** Expression of secIL10 in therapeutic sensor PRB5003 (EcN/pBSIDZ3). Supernatants of PRB5003 induced with 10 and 20 µM TPEN for 2 and 4 h in LB media were collected and used for IL10 ELISA test. Sensor PRB5002 (EcN/pBSIDZ2) with only YebF expression was used as a negative control. Data are mean ± SEM, n = 3 biological independent repeats. Statistical analysis was performed using ANOVA Tukey test; $^{****}p < 0.0001$.

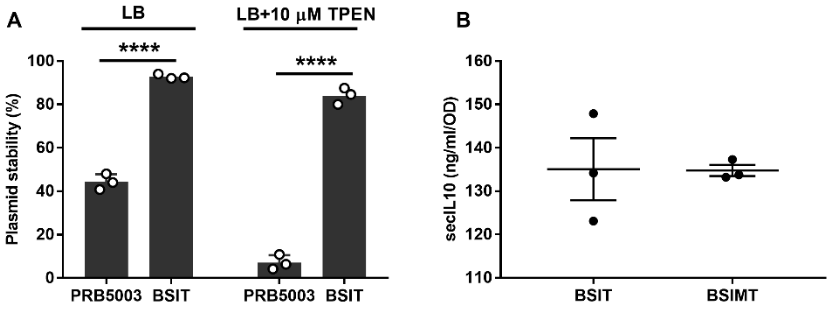


**fig. S5 Plasmid stability and secIL10 expression in stable therapeutic sensor.**

**A:** Plasmid stability of *asd* engineered biosensors without antibiotic selection. Therapeutic biosensors PRB5003 (EcN/pBSIDZ3) and BSIT (Δ*asd*/pBSIT) were cultured in LB media or LB+10 µM TPEN without antibiotic selection, respectively. The biosensors were passaged five times, following 100 µl of 10-fold serial dilution cultures were platted on LB agar plates, then a single colony was replica-cultured in LB and LB with 15 µg/ml chloramphenicol agar plates to calculate plasmid stability. Data are mean ± SEM, n = 3 biological independent repeats. Statistical analysis was performed using an unpaired two-tailed t test; ****$P < 0.0001$.

**B:** Secretion of secIL10 in biosensor BSIT and BSIMT**.** Biosensors were cultured to $OD_{600}$ of 0.5 - 0.6 and then induced with 20 µM TPEN for 4 h in LB media. The supernatants of cultures were collected and detected by IL10 ELISA. Note that expression of IL-10 is similar to BSI1 and BSIM, indicating no impact on IL-10 expression with the addition of asd to the plasmid. Data are mean ± SEM, n = 3 biological independent repeats.

### Supplementary tables

**Table S1. Bacteria and plasmids utilized in this study**

| Strains or plasmids | Genotype and function | Reference |
|---|---|---|
| **Strains** | | |
| BSI | Biosensor EcN harboring pBSI2 | This work |
| BSIM | Memory circuit biosensor EcN harboring pBSIM1 and pBSIM2 | This work |
| BSIT | EcNΔ*asd* harboring pBSIT | This work |
| BSIC | EcNΔ*asd* harboring pBSIC | This work |
| BSIMT | EcNΔ*asd* harboring pBSIMT and pBSIM1 | This work |
| BSIMC | EcNΔ*asd* harboring pBSIMC and pBSIM1 | This work |
| PRB5000 | Biosensor EcN harboring pBSI1 | This work |
| PRB5001 | Biosensor EcN harboring pBSIDZ1 | This work |
| PRB5002 | Biosensor EcN harboring pBSIDZ2 | This work |
| PRB5003 | Biosensor EcN harboring pBSIDZ3 | This work |
| *E. coli* DH5α | Cloning host | Lab stock |
| *Escherichia coli* Nissle 1917 (EcN) | Engineering strain | Lab stock |
| EcNΔ*asd* | EcN deleted *asd* essential gene | This work |
| *C. difficile* R20291 | Ribotype 027 strain, used for biosensor test in a CDI model | Lab stock |
| **Plasmids** | | |
| pBSI2 | pColE1 containing $P_{ykg}$-*sfgfp* and J23109-*zur* construct | This work |
| pBSIM1 | pColE1 containing $P_{ykg}$-*int8* construct | This work |
| pBSIM2 | p15A containing reversed *sfgfp* flanked by integrase 8 recognization sites *attB* and *attP* | This work |
| pBSIT | pColE1 containing $P_{ykg}$-*secIL10*-$P_{asd}$-*asd* construct | This work |
| pBSIC | pColE1 containing $P_{ykg}$-*yebF*-$P_{asd}$-*asd* construct | This work |
| pBSIMT | p15A containing reversed *secIL10* flanked by integrase 8 recognization sites *attB* and *attP* and $P_{asd}$-*asd* construct | This work |
| pBSIMC | p15A containing reversed *yebF* flanked by integrase 8 recognization sites *attB* and *attP* and $P_{asd}$-*asd* construct | This work |
| pBSI1 | pColE1 containing $P_{ykg}$-*sfgfp* construct | This work |
| pBSIDZ1 | pColE1 containing $P_{ykg}$-*SP4-IL10* construct | This work |
| pBSIDZ2 | pColE1 containing $P_{ykg}$-*yebF* construct | This work |
| pBSIDZ3 | pColE1 containing $P_{ykg}$-*secIL10* (*yebF-IL10*) construct | This work |
| pCas | plasmid with Cas9 gene | (*28*) |
| pTargetF | Plasmid used for sgRNA construction for *asd* gene deletion | (*28*) |

**Table S2. Primers utilized in this study**

| Primer | Sequence (5’ to 3’) | PCR product |
|---|---|---|
| 1-F | AGGCCCTTTCGTCTTCACCTCGAGTAACGGCAATAAACTGTTCACTTCAG | $P_{ykg}$ for pBSI1 |
| 1-R | CTAGTATTTCTCCTCTTTCATTTTTACCTGTTATGTTATAACA | |
| 2-F | TGAAAGAGGAGAAATACTAGATGCGTAAAGGCGAAGAGCTGTTCA | *sfgfp* for pBSI1 |
| 2-R | ATGCCTGGTCTAGATTATTATCATCATTTGTACAGTTCATCCATA | |
| 3-F | GCCTTTTTAGTTAGAAAGCTTTAGCGAGGTTTCTTTTTCACCT | *zur* for pBSI2 |
| 3-R | GAATTCAGGAGATAATAT ATGGAAAAGACCACAACGCAGGAGT | |
| 4-F | GCTAGCACAGTCCCTAGGACTGAGCTAGCTGTAAA | J23109 for pBSI2 |
| 4-R | ACGTCTCATTTTCGCCAGATATCGATTTACAGCTAGCTCAGTCCTAGGGACTGTGCTAGC | |
| 5-F | TAAGGATGATTAATAATCTAGACCAGGCATCAAATAAAACGAAAGGCTCA | Linear pBSI1 without *sfgfp* for pBSIM1 |
| 5-R | AATCCTAGTTATTGTTTTGCTAGCCTTTACCTGTTATGTTATAACATAACC | |
| 6-F | AGGCTAGCAAAACAATAACTAGGATTCGAATGAAAGTTGCCGTTTATTGTCGTG | *int8* for pBSIM1 |
| 6-R | ATGCCTGGTCTAGATTATTAATCATCCTTAGCGAAAGCTAAGGA | |
| 7-F | TTGACAGCTAGCTCAGTCCTAGGTATAATGCTAGC | J23119 for pBSIM2 |
| 7-R | GCTAGCATTATACCTAGGACTGAGCTAGCTGTCAA | |
| 8-F | TTGACAGCTAGCTCAGTCCTAGGTATAATGCTAGCAGTTCGATGAGAGCGATAACCAATCATCAGATAACTATGG | *attB-sfgfp-attP* for pBSIM2 |
| 8-R | TCGGGTGGGCCTTTCTGCGTTTATATTAATAAACTATGGAAGTATGTACA | |
| 9-F | AAACAATAACTAGGATTCGAATGAAAAAAATAACCGCGGCGG | *SP4-IL10* for pBSIDZ1 |
| 9-R | TTATTAATCATCCTTAGCGAAAGCTAAGGATTTTTTTATCTGTTAGTTGCGAATTTTCATGGT | |
| 10/11-F | TAGCAAAACAATAACTAGGATTCGAGGAGAAAAACATGAAAAAAAGAGGGGCGTTTTTAG | *yebF* for pBSIDZ2 |
| 10-R | GAAAGCTAAGGATTTTTTTATCTGTTAACGCCGCTGATATTCCGCCATTCCC | |
| 11-R | CGTGCCTTGGCCCGGGCTACGCCGCTGATATTCCGCCA | *yebF* for pBSIDZ3 |
| 12-F | AGCCCGGGCCAAGGCACGCA | *IL10* for *yebF-IL10* fusion expression |
| 12-R | TAAGGATTTTTTTATCTGTTAGTTGCGAATTTTCATGGTCATATACGCT | |
| 13-F | CAGATAAAAAAAATCCTTAGCTTTC | Linear pBSI2 without *sfgfp* for pBSIDZ3 |
| 13-R | TCGAATCCTAGTTATTGTTTTGCTA | |
| 14-F | GGATCCATAATCAGGATCAATAA | $P_{asd}$-*asd* |
| 14-R | TTACGCCAGTTGACGAAGCATCCGA | |
| 15-F | TGCTTCGTCAACTGGCGTAACAGATAAAAAAAATCCTTAGCTTTC | Linear pBSIDZ2 for pBSIC |
| 15-R | TTGATCCTGATTATGGATCC ACGCCGCTGATATTCCGCCATT | |
| 16-F | TGATCCTGATTATGGATCCTTAGTTGCGAATTTTCATGGTCA | Linear pBSIDZ3 for pBSIT |
| 16-R | TTGATCCTGATTATGGATCC ACGCCGCTGATATTCCGCCATT | |
| 17-F | TGCGCTCGGTCGTTCGGCTGCGGCGAGCGGTATCAGCTCACTCAA | For *attB-rsecIL10-attP* and *attB-ryebF-attP* amplification |
| 17-R | TCTGTTGTTTGTCGGTGAACGCTCTCTACTAGAGTCACACTGGCTCACCTTCG | |
| 18-F | GAGAGCGTTCACCGACAAACAACAGA | |

| | | |
|---|---|---|
| 18-R | CGCCGCAGCCGAACGACCGAGCGCAG | Linear pMSIM1 for pBSIMT and pBSIMC |
| T-F | GGTATCAGCTCACTCAAAGG | *sfgfp* flipping detection primers |
| T-R | TGACATCACCATCCAGTTCC | |
| *amp*-Q-F | GAAGATCAGTTGGGTGCACG | Reference gene for *sfgfp* flipping detection |
| *amp*-Q-R | TCACTCATGGTTATGGCAGC | |
| *sfgfp*-Q-F | TGCGGTTTACCAGGGTATCG | *sfgfp* transcription analysis |
| *sfgfp*-Q-R | TATGTGCAGGAACGCACGAT | |
| 16S-F | TAATACCTTTGCTCATTG | Reference gene for transcription analysis |
| 16S-R | CCAGTAATTCCGATTAAC | |

### References


1. A. Cubillos-Ruiz *et al.*, Engineering living therapeutics with synthetic biology. *Nat Rev Drug Discov* **20**, 941-960 (2021).
2. M. R. Charbonneau, V. M. Isabella, N. Li, C. B. Kurtz, Developing a new class of engineered live bacterial therapeutics to treat human diseases. *Nat Commun* **11**, 1738 (2020).
3. B. P. Landry, J. J. Tabor, Engineering Diagnostic and Therapeutic Gut Bacteria. *Microbiol Spectr* **5**, (2017).
4. D. T. Riglar, P. A. Silver, Engineering bacteria for diagnostic and therapeutic applications. *Nat Rev Microbiol* **16**, 214-225 (2018).
5. N. Aggarwal, A. M. E. Breedon, C. M. Davis, I. Y. Hwang, M. W. Chang, Engineering probiotics for therapeutic applications: recent examples and translational outlook. *Curr Opin Biotechnol* **65**, 171-179 (2020).
6. K. N. Daeffler *et al.*, Engineering bacterial thiosulfate and tetrathionate sensors for detecting gut inflammation. *Mol Syst Biol* **13**, 923 (2017).
7. T. Chien *et al.*, Enhancing the tropism of bacteria via genetically programmed biosensors. *Nat Biomed Eng* **6**, 94-104 (2022).
8. L. Liu *et al.*, An Electrochemical Biosensor with Dual Signal Outputs: Toward Simultaneous Quantification of pH and O(2) in the Brain upon Ischemia and in a Tumor during Cancer Starvation Therapy. *Angew Chem Int Ed Engl* **56**, 10471-10475 (2017).
9. N. E. Walsham, R. A. Sherwood, Fecal calprotectin in inflammatory bowel disease. *Clin Exp Gastroenterol* **9**, 21-29 (2016).
10. A. Ricciuto, A. M. Griffiths, Clinical value of fecal calprotectin. *Crit Rev Clin Lab Sci* **56**, 307-320 (2019).
11. T. Vogl, N. Leukert, K. Barczyk, K. Strupat, J. Roth, Biophysical characterization of S100A8 and S100A9 in the absence and presence of bivalent cations. *Biochim Biophys Acta* **1763**, 1298-1306 (2006).
12. M. Fagerhol, K. Andersson, C. Naess-Andresen, P. Brandtzaeg, I. J. B. R. Dale, Fla: CRC Press, Inc, Calprotectin (the L1 leukocyte protein) In: Smith VL, Dedman JR, editors. Stimulus response coupling: the role of intracellular calcium-binding proteins. 187-210 (1990).
13. M. B. Brophy, J. A. Hayden, E. M. Nolan, Calcium ion gradients modulate the zinc affinity and antibacterial activity of human calprotectin. *J Am Chem Soc* **134**, 18089-18100 (2012).
14. T. G. Nakashige, B. Zhang, C. Krebs, E. M. J. N. c. b. Nolan, Human calprotectin is an iron-sequestering host-defense protein. **11**, 765-771 (2015).
15. T. G. Nakashige, E. M. Zygiel, C. L. Drennan, E. M. Nolan, Nickel Sequestration by the Host-Defense Protein Human Calprotectin. *J Am Chem Soc* **139**, 8828-8836 (2017).
16. M. Schultz, Clinical use of E. coli Nissle 1917 in inflammatory bowel disease. *Inflamm Bowel Dis* **14**, 1012-1018 (2008).
17. J. P. Lynch, L. Goers, C. F. Lesser, Emerging strategies for engineering Escherichia coli Nissle 1917-based therapeutics. *Trends Pharmacol Sci* **43**, 772-786 (2022).
18. G. Zhang, S. Brokx, J. H. Weiner, Extracellular accumulation of recombinant proteins fused to the carrier protein YebF in Escherichia coli. *Nat Biotechnol* **24**, 100-104 (2006).
19. S. E. Gabriel, J. D. Helmann, Contributions of Zur-controlled ribosomal proteins to growth under zinc starvation conditions. *J Bacteriol* **191**, 6116-6122 (2009).

20. J. Z. Liu *et al.*, Zinc sequestration by the neutrophil protein calprotectin enhances Salmonella growth in the inflamed gut. *Cell Host Microbe* **11**, 227-239 (2012).
21. C. McDowell, M. Haseeb, Inflammatory bowel disease (IBD). (2017).
22. S. Jose, R. Madan, Neutrophil-mediated inflammation in the pathogenesis of Clostridium difficile infections. *Anaerobe* **41**, 85-90 (2016).
23. A. M. Buckley, J. Spencer, D. Candlish, J. J. Irvine, G. R. Douce, Infection of hamsters with the UK Clostridium difficile ribotype 027 outbreak strain R20291. *J Med Microbiol* **60**, 1174-1180 (2011).
24. D. Zhu, J. Bullock, Y. He, X. Sun, Cwp22, a novel peptidoglycan cross-linking enzyme, plays pleiotropic roles in Clostridioides difficile. *Environ Microbiol* **21**, 3076-3090 (2019).
25. E. M. Panina, A. A. Mironov, M. S. Gelfand, Comparative genomics of bacterial zinc regulons: enhanced ion transport, pathogenesis, and rearrangement of ribosomal proteins. *Proc Natl Acad Sci U S A* **100**, 9912-9917 (2003).
26. D. Kandari, H. Joshi, R. Bhatnagar, Zur: Zinc-Sensing Transcriptional Regulator in a Diverse Set of Bacterial Species. *Pathogens* **10**, (2021).
27. J. Perez, V. S. Springthorpe, S. A. Sattar, Clospore: a liquid medium for producing high titers of semi-purified spores of Clostridium difficile. *J AOAC Int* **94**, 618-626 (2011).
28. Y. Jiang *et al.*, Multigene editing in the Escherichia coli genome via the CRISPR-Cas9 system. *Appl Environ Microbiol* **81**, 2506-2514 (2015).
